# Quantification of cannabidiolic acid in *Cannabis sativa* using a two-component allosteric protein biosensor

**DOI:** 10.64898/2026.11.17.767439

**Authors:** Zhong Guo, Carlos Luna-Flores, Raine E. S. Thomson, Cade Turland, David Grolman, Kirill Alexandrov, James B. Behrendorff

## Abstract

Cannabidiol (CBD) is an important medicinal cannabinoid which accumulates in the *Cannabis sativa* plant as the precursor cannabidiolic acid (CBDA). Measurement of CBDA levels is important for crop valuation, harvest optimisation, and quality control. However, its quantification typically requires laboratory-based analytical methods that are not readily deployable at the point of need. Here, we report the development of an engineered protein biosensor for selective CBDA detection based on chemically induced dimerization of engineered nanobody domains. Here a cannabidiol-responsive nanobody pair was incorporated into a two-component β-lactamase-based biosensor architecture, such that ligand binding ultimately results in reconstitution of β-lactamase activity quantified in a simple colorimetric assay. The resulting sensor displayed high affinity for CBD (*K*_*D*_ = 1.9 ± 0.2 nM) and CBDA (*K*_*D*_ = 32 ± 4 nM). Structure analysis informed a single substitution mutation of ligand-binding nanobody to shift its selectivity in favour of CBDA. This minimal modification inverted sensor specificity, generating a CBDA-selective biosensor that showed no detectable activation either by CBD or the immediate biosynthetic precursor, cannabigerolic acid (CBGA). The engineered sensor exhibited nanomolar affinity for CBDA (*K*_*D*_ = 21 ± 2.4 nM), an 8.4-fold dynamic range, and a limit of detection of 0.5 nM. Sensor performance was validated using crude extracts from CBDA-producing biomass and achieved 95.6% agreement with LCMS-based quantification.

## Introduction

Medicinal crops including *Cannabis sativa* are cultivated for high-value secondary metabolites. Monitoring the value-determining maturity of medicinal crops requires sophisticated analytical chemistry that cannot be deployed in the field (Harpaz *et al*. 2022). Growers are often reliant on third-party analytical services that can be time-consuming and costly, limiting the spatial and temporal resolution of crop data. At current per-sample costs, cannabinoid content is commonly measured only to meet regulatory requirements (Berthold *et al*. 2020, Arsenault *et al*. 2024). For example, in Australian medicinal cannabis and hemp production, a single cannabinoid potency assay performed by an accredited third-party laboratory costs approximately AUD $176 per sample with a typical turnaround of 7-10 business days. A licensed cultivation site may commission dozens of such assays per crop cycle across cultivar screening, harvest timing, in-process checks and batch release to ensure quality and compliance. Cannabis cultivation methods and harvest timing are decided largely by visual assessment of plants and calendar heuristics rather than measured cannabinoid accumulation, in part due to the associated cost of cannabinoid quantification. However, changes in temperature, day length, planting density, and fertigation can have a substantial impact on cannabinoid yield (Ii *et al*. 2021, De Prato *et al*. 2022, Malík *et al*. 2025). Low cost, on-farm methods for detection and measurement of specific cannabinoids would create new opportunities for optimising growth conditions and harvest time. Highly specific detection methods that can discriminate between structurally similar chemical compounds would also enable efficient identification of cultivars that are enriched in preferred cannabinoids.

Emerging technologies for rapid and portable cannabinoid detection include lateral flow immunoassays, electrochemical sensors, and engineered live cell biosensors (Plouffe and Murthy 2017, Chand *et al*. 2021, Amini *et al*. 2022, Miettinen *et al*. 2022, Mulvihill *et al*. 2024). Electrochemical and immunoassay technologies for detecting cannabinoids are typically limited by their sensitivity to matrix impurities and non-target analytes. Engineered live cell sensors can have high tolerance for matrix impurities and high target specificity, but their deployment in the field is often impractical due to regulations surrounding release of genetically modified organisms as well as to limitations related to typically slow response times of sensors that depend on transcriptional activation and accumulation of a measurable protein product. Furthermore, existing live cell cannabinoid sensor devices are mostly designed for detection of decarboxylated cannabinoid end products rather than the carboxylate form that accumulates *in planta*, and this limits their on-farm utility.

Protein biosensors consist of a ligand-binding receptor component(s) that is engineered for high affinity and specificity towards the target analyte, and a reporter component that converts binding events into a measurable signal such as fluorescence or enzymatic activity (Guo *et al*. 2026). These molecular devices can provide the same benefits as cell-based sensors including exquisite molecular selectivity, quantitative analysis, and tolerance of matrix impurities, but without the practical complications of maintaining live microorganisms (Jackson, Anderson, and Alexandrov 2022).

Previously a chemically-inducible protein dimerization system specific for cannabidiol (CBD) was developed using a pair of protein nanobody domains selected from a phage display library (Kang *et al*. 2019, Cao *et al*. 2022). We adapted these CBD binding domains to an existing two-component protein biosensor architecture where chemically induced dimerization triggers activation of β-lactamase reporter domain, such that β-lactamase enzyme activity is proportional to analyte concentration (**Figure 1**) (Guo *et al*. 2023). Here we report functional validation of this CBD-specific two component biosensor and undertake further protein engineering to change the specificity of the ligand binding domains from CBD to cannabidiolic acid (CBDA). Finally, we validate performance of the CBDA biosensor using crude biomass extracts from a CBDA-rich hemp variety.

**Figure 1.**
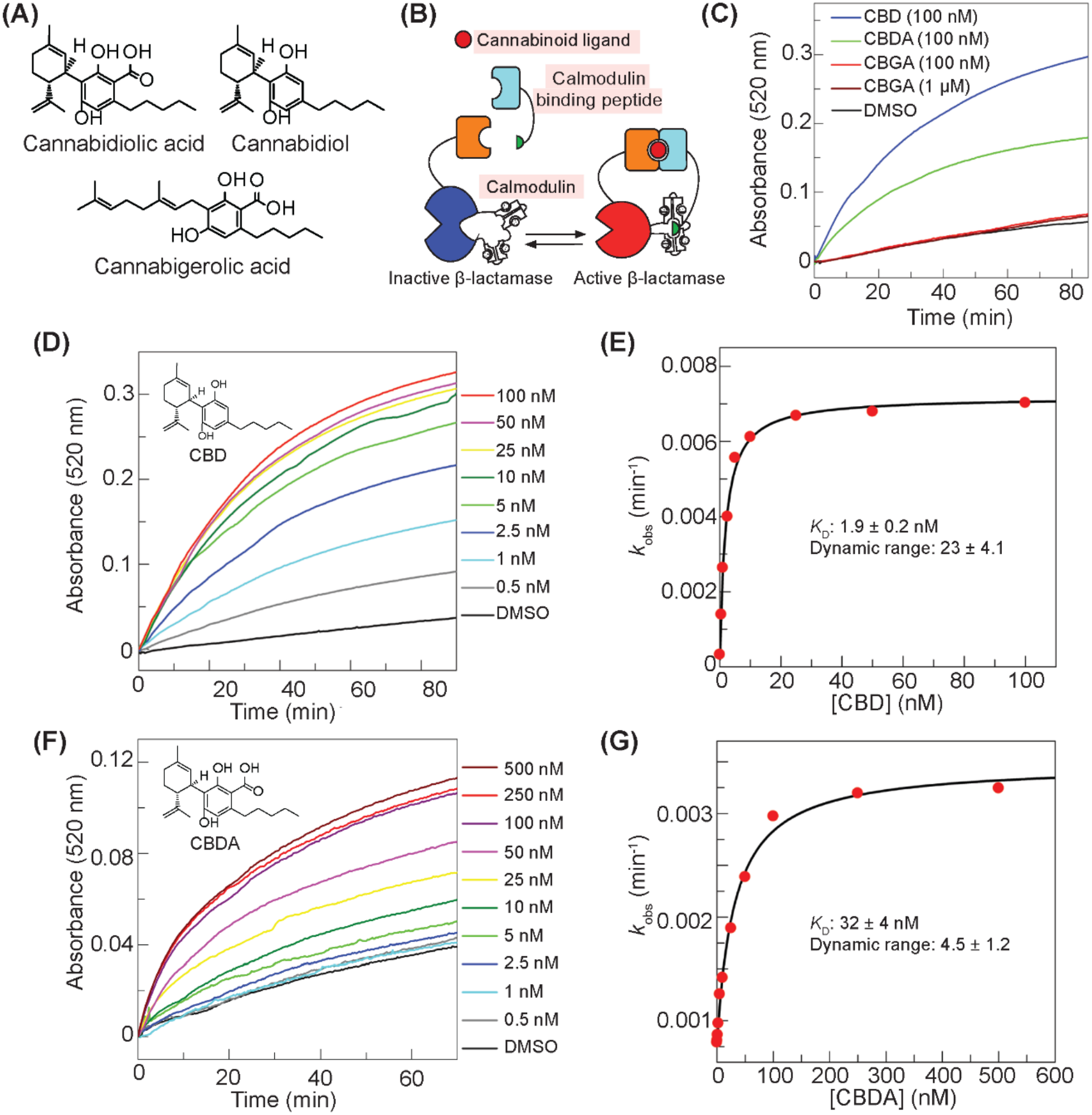
Architecture and activity analysis of two-component biosensor of cannabidiolic acid and related compounds. **(A)** Chemical structures of cannabidiolic acid (CBDA), the decarboxylated product cannabidiol (CBD), and the metabolic precursor cannabigerolic acid (CBGA). **(B)** Schematic representation of the two-component biosensor mechanism, where binding of a cannabinoid ligand recruits the calmodulin-binding peptide (CA14-CaM BP) to calmodulin, causing allosteric activation of the β-lactamase reporter domain (CaM-BLA-DB21). **(C)** Plot showing change in absorbance at 520 nm due to β-lactamase-mediated hydrolysis of UW154 (50 µM) upon incubation of the two-component biosensor (10 nM CaM-BLA-DB21, 100 nM CA14-CaM BP) with 100 nM CBD, CBDA, CBGA, or 1 µM CBGA at 25 °C. **(D)** β-lactamase activity of a 10 nM solution of CaM-BLA-DB21 and 100 nM CA14-CaM BP in the presence of the increasing concentrations of CBD (0-100 nM) analysed by changes in UW154 absorbance. **(E)** Initial rate (*k*_obs_, min^−1^) of UW154 hydrolysis in response to different CBD concentrations (0-100 nM) from **(D)** fitted to a quadratic binding equation to determine dissociation constant (*K*_D_) and the dynamic range. **(F)** β-lactamase activity of a 10 nM solution of CaM-BLA-DB21 and 100 nM CA14-CaM BP in the presence of the increasing concentrations of CBDA (0-500 nM) analysed by changes in UW154 absorbance. **(G)** Initial rate (*k*_obs_, min^−1^) of UW154 hydrolysis in response to different CBDA concentrations (0-500 nM) from **(F)** fitted to a quadratic binding equation to determine dissociation constant (*K*_D_) and the dynamic range. Calculated values presented in (E) and (G) represent mean ± SD from *n* = 3 independent replicates.

## Materials and Methods

### Cannabinoid preparation and analytical chemistry

Cannabinoid analytical standards were purchased from Supelco (Novachem Australia catalogue numbers: CBD, cat no. C-045-1ML; CBDA, cat no. C-144-1ML). Dried hemp biomass rich in either CBDA or CBGA was provided by Hale Farm Pty Ltd (Australia). Cannabinoid extracts were prepared from 1 g dry sifted hemp biomass via dynamic maceration in 10 mL of 100% ethanol (Brighenti *et al*. 2017), stirring at room temperature for 15 min. Insoluble material was sedimented via centrifugation at 3,500 *g* for 5 min, and the resulting supernatant was collected. The residual biomass was re-extracted three further times in 5 mL of fresh ethanol. Supernatants were combined and filtered through a 0.45 μm filter. The pooled extract was dried under a continuous stream of nitrogen and stored at –80 °C.

Dried biomass extracts were re-dissolved in 100% dimethyl sulfoxide (DMSO). Prior to analysis, analytical standards and biomass extracts were diluted in a solution of 60% acetonitrile with 0.1% formic acid. CBD and CBDA content were analysed via LC-MS/MS with a Shimadzu LCMS-8050 instrument equipped with a DUIS dual ion source, coupled to a Nextera UPLC system (Shimadzu). Samples (0.2 µL injection volume) were applied to a 5 µm C18 100 Å column (150 × 4.6 mm, Phenomenex cat no. 00F-4633-E0). The mobile phase consisted of 0.1% formic acid in water (solvent A) and 0.1% formic acid in acetonitrile (solvent B). Samples were eluted in a mobile phase that increased from 60% solvent B to 80% solvent B in a linear gradient over 15 minutes, followed by re-equilibration to 60% solvent B from 15-20 minutes. Mass spectrometry analysis was conducted in negative ion mode for CBDA and positive mode for CBD, targeting specific mass-to-charge ratios (m/z) for cannabinoid detection. The transitions monitored for cannabidiolic acid (CBDA, retention time 5.8 min) were *m/z* 357 → 245 (collision energy 29 eV), 357 → 339 (collision energy 21 eV), and 357 → 179 (collision energy 26 eV). For cannabidiol (CBD, retention time 6.4 min), the transitions monitored were 315 → 123 (collision energy −35 eV), 315 → 81 (collision energy −18 eV), 315 → 259 (collision energy −19 eV), and 315 > 193 (collision energy −21 eV). Data analysis and quantification were performed using Skyline-daily (64-bit) version 23.1.1.425, with CBDA and CBD standards serving as references for calibration and quantitation.

### Molecular cloning, expression, and purification of recombinant proteins

The open reading frames of the constructs listed in Supplementary Table S1 were synthesized commercially (Gene Universal) and cloned into a kanamycin-resistant pET-28a(+) vector. Competent *E. coli* BL21(DE3) cells (New England Biolabs) were transformed with the resulting expression vectors and grown in LB broth with 50 μg/mL kanamycin (Everest Inc, Australia) with shaking at 37 °C. Protein expression was induced by adding 0.3 mM IPTG (Sigma, USA) and the cultures were incubated overnight at 18 °C. Cells were harvested at 4,000 rpm for 10 min, and the cell pellet was lysed in buffer containing 50 mM Na_2_HPO_4_ (pH 8.0), 300 mM NaCl, 20 mM imidazole, 1 mM AEBSF and DNAse I (all Sigma, USA). Following cell lysis and disruption at 27 kPsi using the CFII cell disrupter (Constant Systems, UK), the supernatant was collected, and recombinant protein was further purified on the Ni-NTA HisTrap FF crude column driven by ÄKTA Express Purifier (Cytiva) system in buffer containing 50 mM Na_2_HPO_4_, 300 mM NaCl and 20 mM imidazole and a gradient of 500 mM imidazole for elution at pH 8.0. Finally, the protein was dialyzed against 20 mM Tris/HCl (pH 7.0) and 100 mM NaCl and protein aliquots were frozen in liquid nitrogen and stored at −80 °C.

### Spectrophotometric analysis of β-lactamase enzymatic activity

Enzyme assays were performed in an assay buffer containing 20 mM Tris-Cl (pH 7.2), 100 mM NaCl and 1 mM CaCl_2_ at 25 °C, and monitored the increase in absorbance caused by hydrolysis of 50 µM UW154 (Ghavami *et al*. 2015) at 520 nm using a Cary 60 UV-VIS absorbance spectrometer operated by Cary WinUV Software.

### Quantification of biosensor performance parameters and data plotting

To determine the dynamic range of the individual sensor constructs, we used ligand titration experiments to establish the *K*_*D*_ value for the ligand. We then performed the assay either at zero or saturating concentration of the ligand (typically 20-fold greater than the *K*_*d*_ value) and recorded the absorbance change over time.

The linear phase of the curves recorded at different concentrations of the ligand was fitted as a linear function to obtain the observed rate constant, *k*_obs_. The dynamic range was calculated by dividing the *k*_obs_ of the background signal in the absence of the ligand by the *k*_obs_ recorded at saturating ligand concentration. The mean *k*_obs_ was obtained by averaging three independent experimental data. In order to obtain the *K*_*D*_ for the interaction of biosensors with their ligand, the *k*_obs_ data were plotted against the concentration of the ligand and the data were fitted to the explicit solution of the quadratic equation describing the E + S <> ES binding equilibrium, where *K*_*d*_ is defined as *K*_*D*_= [E]*[S]/[EL]. [E_0_] and [L_0_] refer to the total enzyme and ligand concentration (free and bound) in the cuvette. Under these conditions the *k*_obs_ is described by Equation 1:

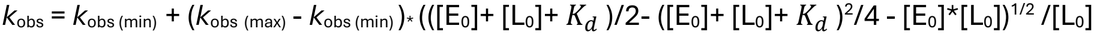

*k*_obs_ represents the measured rate, while k_obs (min)_ and k_obs (max)_ refer to the minimal and maximal rates observed, respectively. A least-squares fit of the data to Equation 1 using the software package Grafit 5.04 (Erithacus software) was used to extract the *K*_*D*_ value and its standard deviation.

## Results

### Development of cannabinoid-controlled two-component biosensor based on β-lactamase

In our previous study, we developed a TEM-1 β-lactamase-based two-component biosensor driven by a chemically induced protein dimerization (CID) system for the therapeutic drug methotrexate (MTX) (Guo *et al*. 2023). As illustrated in **Figure 1B**, MTX-binding VHH domain was fused to a calmodulin-β-lactamase (CaM-BLA) chimera, whereas a binding domain that selectively recognises VHH:MTX was fused to a low affinity M13 calmodulin-binding peptide. Ligand binding induces ternary complex formation between the two binders and the target analyte, locally concentrating the low-affinity M13 peptide in the proximity of the chimeric reporter protein. Consequently, binding of the M13 peptide to the calmodulin moiety to CaM-BLA chimera induces it structural rearrangement accompanied by the increase in catalytic activity.

To construct a cannabinoid-responsive two-component biosensor based on β-lactamase, we searched the Protein Data Bank (PDB) for cannabinoid-mediated CID systems. From this structural analysis, we selected a dual-nanobody system that binds cannabidiol (CBD) with high cooperativity (Cao *et al*. 2022). We generated two distinct fusion constructs by coupling each nanobody to either the CaM-BLA chimera or the engineered low-affinity M13 peptide via flexible linkers (protein sequences are provided in **Supplementary Table 1**).

The resulting two-component biosensor pairs—CaM-BLA-CA14 / DB21-CaM BP and CaM-BLA-DB21 / CA14-CaM BP—were successfully expressed in *E*.*coli* and purified to homogeneity by affinity chromatography. Enzymatic activity was evaluated using a colorimetric assay with the chromogenic β-lactamase substrate UW154 (Ghavami *et al*. 2015). Notably, CBD dose-dependently activated the CaM-BLA-DB21 / CA14-CaM BP sensor pair, whereas the inverse CaM-BLA-CA14 / DB21-CaM BP configuration exhibited no detectable activation. These findings indicate that the spatial orientation and domain configuration of the ligand-binding domains are critical for functional sensor activation.

We next investigated the selectivity of the active biosensor (CaM-BLA-DB21 / CA14-CaM BP) against metabolic precursors along the CBD biosynthetic pathway, specifically evaluating CBD, cannabidiolic acid (CBDA), and cannabigerolic acid (CBGA). The biosensor was activated by both CBD and CBDA, whereas CBGA failed to elicit a response (**Figure 1C**). This suggests that the cyclohexene ring that is present in CBD and CBDA, but absent in the acyclic precursor CBGA, is necessary for heterodimerization between the DB21 and CA14 nanobodies.

Subsequent dose-response titrations with increasing concentrations of CBD and CBDA (**Figure 1D, F**) yielded dissociation constants (*K*_*D*_) and dynamic ranges for each analyte (**Figure 1E, G**). The biosensor exhibited a *K*_*D*_ of 1.9 ± 0.2 nM and a dynamic range of 23 ± 4.1 fold for CBD. By comparison, biosensor activation by CBDA binding exhibited a *K*_*d*_ of 32 ± 4 nM and a dynamic range of 4.5 ± 1.2 fold. Thus, the sensor demonstrates approximately 17-fold binding selectivity for CBD over CBDA, with a limit of detection of at least 0.5 nM for CBD in this biochemical format.

Structural analysis of the CA14:DB21:CBD ternary complex crystal structure reveals a molecular basis for this binding hierarchy. The additional carboxyl group (−COOH) present at the 2-position of the aromatic ring in CBDA introduces both steric hindrance and electrostatic repulsion with residue Asp34 (D34) of nanobody CA14 (**Figure 2A**). This localized steric clash and unfavourable charge interaction reduce the binding affinity of the biosensor toward CBDA relative to CBD.

**Figure 2.**
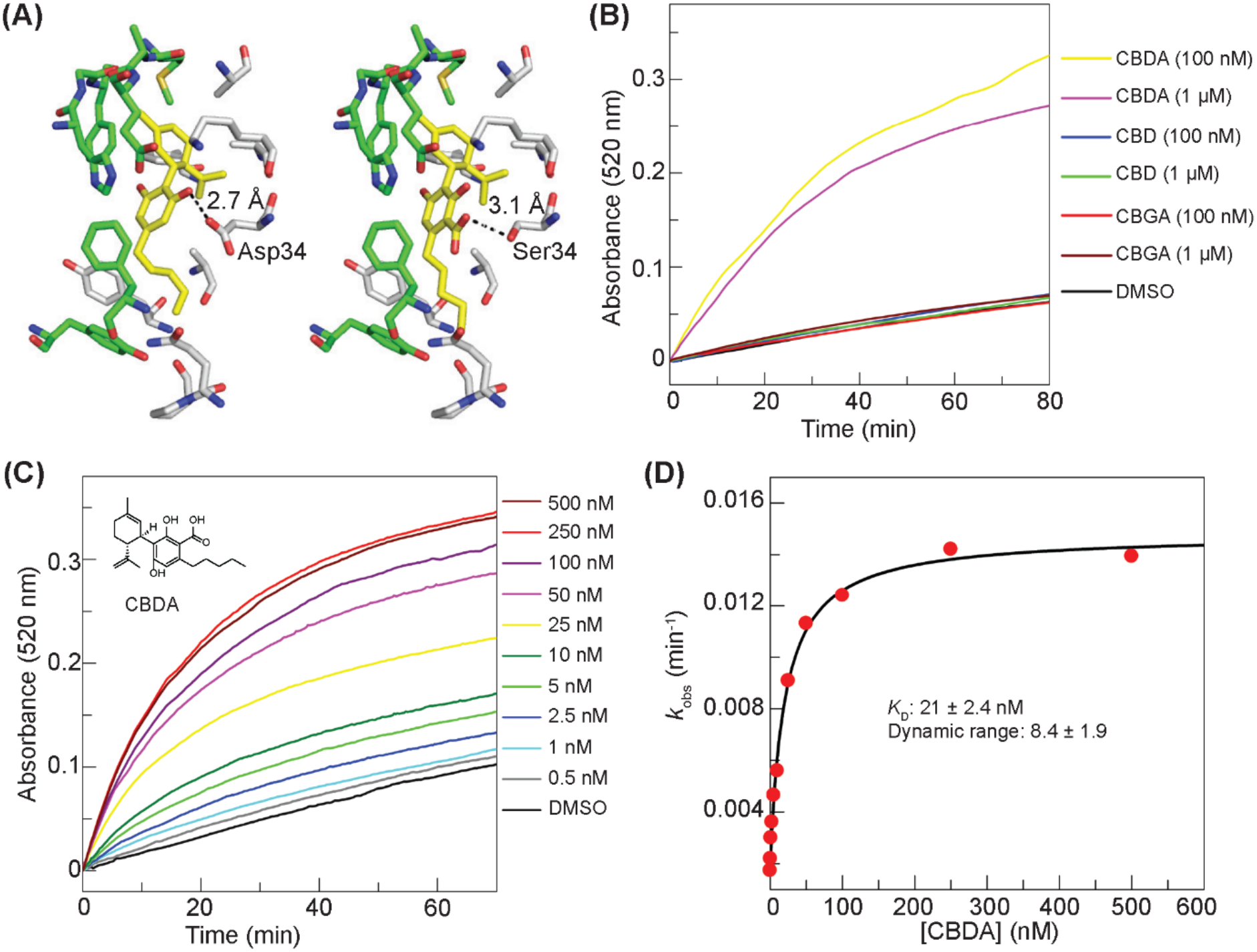
Engineering biosensor specificity toward cannabidiolic acid. **(A)** Structural model of the ligand-binding pocket based on the crystal structure of the CBD-specific DB21:CA14:CBD ternary complex (PDB: 7TE8, left) placed next to the Boltz-1 predicted structural model of the engineered DB21:CA14_D34S_:CBDA complex (right). Amino acid residues within 4 Å of the ligand are depicted. Stick backbones of DB21, CA14, and CBD/CBDA are colored green, grey, and yellow, respectively. **(B)** Change in absorbance at 520 nm due to β-lactamase-mediated hydrolysis of UW154 (50 µM) upon incubation of the two-component biosensor (10 nM CaM-BLA-DB21, 100 nM CA14-CaM BP) in the presence of CBD, CBDA, or CBGA (each at 100 nM or 1 µM) at an incubation temperature 25 °C. **(C)** β-lactamase activity of a 10 nM solution of CaM-BLA-DB21 and 100 nM CA14-CaM BP in the presence of the increasing concentrations of CBDA (0-500 nM) analysed by changes in UW154 absorbance. **(D)** Initial rate (*k*_obs_, min^−1^) of UW154 hydrolysis in response to different CBDA concentrations (0-500 nM) from **(C)** fitted to a quadratic binding equation to determine dissociation constant (*K*_D_) and the dynamic range. Data represent mean ± SD (*n* = 3).

### Engineering biosensor specificity toward cannabidiolic acid

Cannabinoids accumulate *in planta* as carboxylic acids (e.g. CBDA or THCA), which are thermally decarboxylated to the therapeutic cannabinoid end products (e.g. CBD or THC) during post-harvest processing. A biosensor specific for CBDA is needed to enable quantification of this cannabinoid in plant tissue. To adapt the CBD biosensor for CBDA detection, we re-engineered the biosensor’s binding pocket for CBDA specificity through site-directed mutagenesis of residue Asp34 (D34) in nanobody CA14.

To improve specificity toward CBDA, we substituted Asp34 with serine (D34S) based on three rational design strategies: (1) disrupting the 2.7 Å hydrogen bond between Asp34 and the hydroxyl group of CBD, (2) relieving the steric clash with the carboxyl group of CBDA, and (3) introducing a novel hydrogen-bonding interaction between the mutated Ser34 residue and the carboxylic acid group of CBDA. Structural modelling of the CA14(D34S):DB21:CBDA ternary complex using Boltz-1(Wohlwend *et al*. 2025) predicted the formation of a 3.1 Å hydrogen bond between Ser34 and the carboxyl moiety of CBDA (**Figure 2A**).

We expressed and purified the mutated biosensor variant (CaM-BLA-DB21/CA14(D34S)-CaM BP) and evaluated its substrate selectivity. Biochemical assays demonstrated that the engineered biosensor was exclusively activated by CBDA, exhibiting no detectable response to either CBD or CBGA. Dose-response titrations with increasing concentrations of CBDA (**Figure 2C**) yielded an equilibrium dissociation constant (*K*_*D*_) of 21 ± 2.4 nM and a dynamic range of 8.4 ± 1.9 fold (**Figure 2D**). These titration data confirm that the engineered CBDA-specific biosensor achieves a limit of detection down to 0.5 nM for CBDA in this biochemical format.

### Validation of biosensor performance with hemp biomass samples

Sensitivity to off-target hemp metabolites was tested using industrial hemp, which is rich in the cannabinoid precursor CBGA but contains only trace quantities of therapeutic cannabinoids. Ethanolic extracts of dried CBGA-rich hemp biomass were resuspended in DMSO and diluted to a final concentration of 100 nM or 1 µM CBGA. Biosensor response to extracts of CBGA-rich industrial hemp was indistinguishable from negative controls, demonstrating that the biosensor is not activated by non-cannabinoid metabolites present in ethanolic cannabis extracts (**Supplementary Figure S1**).

The performance and practical utility of our CBDA-specific biosensor were validated by evaluating five independent samples extracted from CBDA-rich hemp biomass. Ethanolic extracts of CBDA-rich biomass were resuspended in DMSO and the concentrations of CBDA and CBD were determined via LC-MS/MS (**Supplementary Figure S2**). Biomass extracts quantified via LC-MS/MS were diluted to a final CBDA concentration of 50 nM and incubated with the CBDA-specific biosensor, with comparison to a 50 nM CBDA reference standard. Trace quantities of CBD were also observed in extracts of CBDA-rich hemp (**Supplementary Figure S2**). The biosensor exhibited comparable activation between the pure CBDA standard and hemp extract samples (**Figure 3A**). Furthermore, comparing the observed rate constants (*k*_obs_) of the hemp extract samples against the reference standard yielded a calculated CBDA concentration of 47.8 ± 1.8 nM for the extract samples, demonstrating close agreement with the hemp extract CBDA concentration determined via LC-MS/MS (95.6 % accuracy) **(Figure 3B)**. These results confirm that the engineered biosensor assay provides an accurate and reliable platform for quantifying CBDA concentration and verifying its presence directly within complex plant matrices with minimal interference by co-existing hemp metabolites.

**Figure 3.**
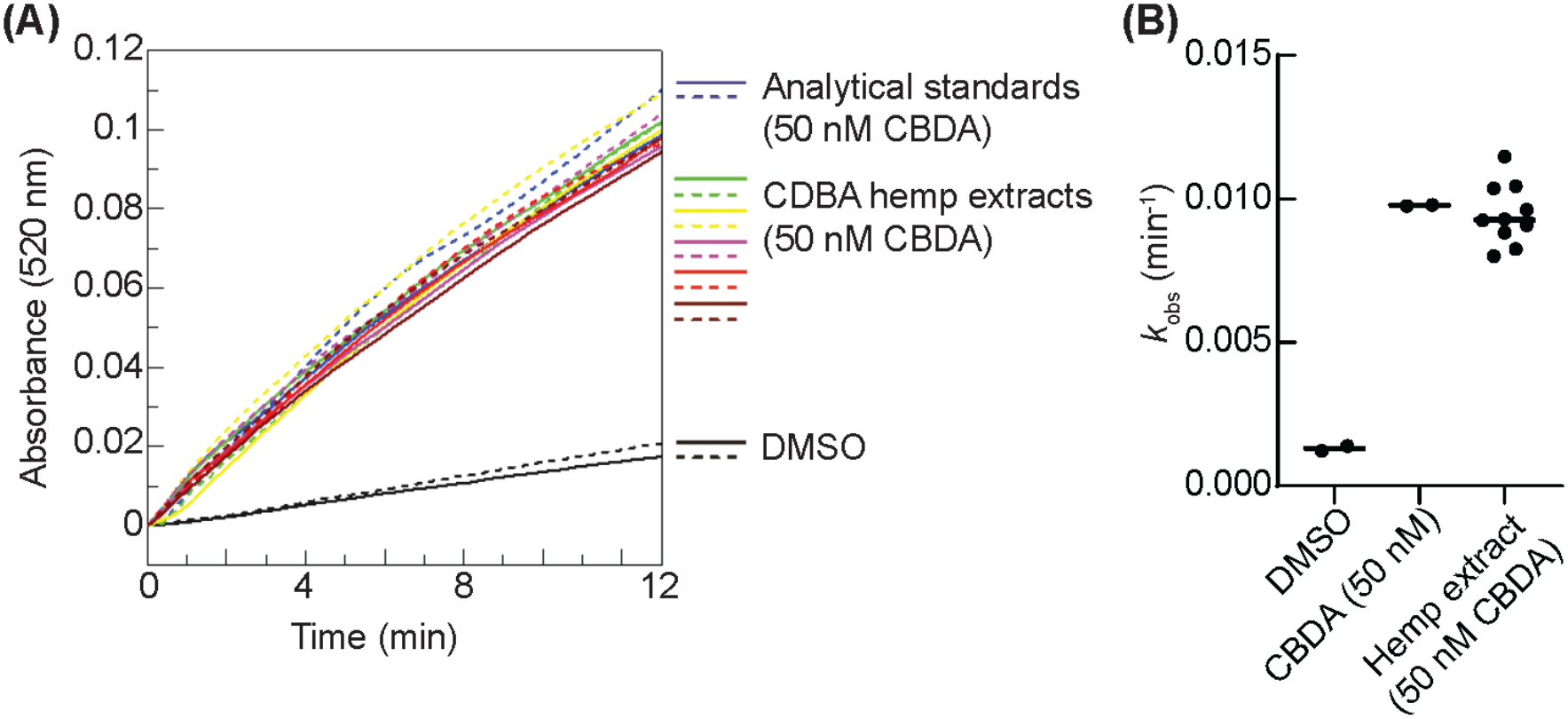
Activation of the engineered CBDA-specific biosensor by CBDA present in cannabis biomass extracts. **(A)** Response of the CBDA biosensor (10 nM CaM-BLA-DB21 and 100 nM CA14_D34S_-CaM BP) to CBDA analytical standards (50 nM, two independent replicates) or CBDA-rich hemp extracts diluted to a CDBA concentration of 50 nM (five independent hemp extracts indicated with different colours, each assayed with two technical replicate incubations indicated by solid and dashed lines). Biosensor-mediated hydrolysis of UW154 (50 µM) is reported as change in absorbance at 520 nm at 25 °C. **(B)** Observed rate constant (*k*_obs_, min^−1^) calculated from **(A)** describing rate of UW154 hydrolysis in the presence of analytical standards or samples of CBDA-hemp extracts each containing 50 µM CBDA.

## Discussion

Chemically-induced dimerization of proteins is a key mechanism of chemical information transduction in biological systems (Stanton, Chory, and Crabtree 2018). Using protein engineering, the specificity of small molecule binding domains can be altered in favour of desired target analytes and combined with reporter domains that convert binding events into readable signals (Foight *et al*. 2019, Jackson, Anderson, and Alexandrov 2022).

In this study, we re-engineered a TEM-1 β-lactamase-based two-component biosensor to achieve strict binding selectivity for cannabidiolic acid (CBDA) over its neutral derivative (CBD) and precursor (CBGA). Initial characterization of a CBD-binding nanobody pair (CA14 and DB21), constructed using sequential phage display of VHH library, revealed a strong preference for CBD (*K*_*D*_ = 1.9 nM) compared to CBDA (*K*_*D*_ = 32 nM). Structural modelling identified a key molecular feature driving this bias: the carboxyl group (−COOH) at the 2-position of the aromatic ring in CBDA introduces significant steric hindrance and unfavourable electrostatic repulsion with residue Asp34 (D34) on nanobody CA14.

To overcome this structural barrier, we implemented a rational single-point mutation (D34S) on CA14. Substituting aspartate with serine improved selectivity through elimination of the 2.7 Å hydrogen bond anchored to the hydroxyl group of CBD, relief of the steric clash with the carboxyl group of CBDA, and by establishing a new 3.1 Å hydrogen-bonding network between Ser34 and the carboxylic acid moiety of CBDA. This minimal structural adaptation completely inverted the sensor’s substrate selectivity profile, yielding an engineered biosensor (CaM-BLA-DB21 / CA14(D34S)-CaM BP) that exhibits high affinity for CBDA (*K*_*D*_ = 21 nM) with no detectable activation by CBD or CBGA.

A central challenge in adapting biosensors for real-world agricultural and industrial applications is matrix interference from crude biological extracts. Industrial hemp biomass contains a complex mixture of non-target terpenes, chlorophylls, and cannabinoid precursors such as CBGA. Previous studies have expressed cannabinoid-responsive G protein-coupled receptors (GPCRs) in *Saccharomyces cerevisiae* and coupled their activation to expression of measurable reporters such as fluorescent proteins or luciferase enzyme. However these sensors are constructed from protein receptors that have evolved as for broad ligand recognition, and interact with a wide range of natural and synthetic cannabinoids, as well as other phytochemicals, precluding their use in determining the presence of specific cannabinoids in a complex plant extract (Miettinen *et al*. 2022, Shaw *et al*. 2022, Mulvihill *et al*. 2024). Our engineered protein biosensor maintained high fidelity when challenged with crude extracts of CBDA-rich cannabis, producing signal outputs directly comparable to those observed with purified CBDA standards. The absence of background signal from CBGA-rich hemp biomass underscores the specificity achieved through the D34S mutation, confirming that the open-chain architecture of CBGA lacks the necessary cyclohexene ring contacts required to stabilize the active heterodimeric state of the biosensor.

Biosensor technologies do not replace accredited laboratory testing required for regulatory compliance, but could support on-site analyses for optimisation of cultivation practices, and crop valuation, as well as validation of third-party biomass (Milay *et al*. 2020, Chen *et al*. 2023). Monitoring CBDA loss through decarboxylation could also enable optimisation of post-harvest processing and storage conditions. Cannabinoid yields can vary substantially among seedlings derived from a single parent plant and even among clonal cuttings; early analysis of seedlings and clonal cuttings may allow for culling of low-productivity plant lines before committing to a full production cycle (Dilena, Hunt, and Close 2025, Garcia-de Heer *et al*. 2026). The protein biosensor technology presented here can potentially be developed for field deployment: the analyte is prepared as a crude biomass extract that can be prepared on-site, and the colorimetric output eliminates the need for specialist laboratory equipment. Changes in absorbance can be readily monitored using accessible, portable, and low-cost colorimetric hardware. For example, ambient colorimetric rate measurements can be recorded using consumer smartphones equipped with basic camera-based RGB-analysing algorithms, while designs for self-assembled single-wavelength LED photometers are widely available online (Wang *et al*. 2015, Li *et al*. 2024, Tang *et al*. 2025). This bridges the gap between high-precision protein engineering and low-cost, point-of-need analytical testing, enabling direct monitoring of CBDA content in cannabis biomass.

## Conclusion

We developed and rationally engineered a highly selective, two-component β-lactamase biosensor for the direct detection and quantification of cannabidiolic acid (CBDA). Initial evaluation of a CBD-specific nanobody pair (CA14 and DB21) demonstrated a strong preference for CBD over CBDA, driven by steric hindrance and electrostatic repulsion between residue Asp34 (D34) on nanobody CA14 and the carboxyl moiety of CBDA. Guided by structural modelling, we implemented a rational single-point mutation (CA14_D34S_) that inverted this target preference, eliminating cross-reactivity toward both CBD and the acyclic precursor CBGA while achieving a nanomolar dissociation constant *K*_*D*_ = 21 2.4 nM (*K*_*D*_ = 21 ± 2.4 nM) and a limit of detection down to 0.5 nM for CBDA.

## Author contributions

ZG developed, engineered, and characterised the CBD and CBDA biosensors. CLF and REST contributed cannabinoid extraction and analytical chemistry. CT and DG prepared and contributed cannabis biomass. ZG, KA, and JBB designed the study and analysed the data. All authors contributed to preparation of the manuscript.

## Acknowledgements

This work was supported in part by the Australian Research Council Centre of Excellence in Synthetic Biology (project CE200100029, awarded to K.A, R. E. S., and J. B. B.), Australia’s Economic Accelerator (project AE230200068, awarded to C. L. F., J. B. B., D. G., and C. T.), ARC Linkage Project LP200200916 (awarded to K. A.), and NHRMC Investigator grant APP2033951 (awarded to K. A.), and in-kind contributions of hemp biomass from Hale Farm Pty Ltd. K.A. gratefully acknowledges financial support from the CSIRO-QUT Synthetic Biology Alliance and the CSIRO Synthetic Biology and Advanced Engineering Biology Future Science Platforms. Analytical chemistry was performed at the Central Analytical Research Facility operated by QUT Research Infrastructure.

## Data availability

The data underlying this article will be shared on reasonable request to the corresponding author.

## References

Amini K, Sepehrifard A, Valinasabpouri A et al. Recent advances in electrochemical sensor technologies for THC detection—a narrative review. J Cannabis Res 2022;4(4):12. 10.1186/s42238-022-00122-3.

Arsenault TL, Prapayotin-Riveros K, Ammirata MA et al. Compliance Testing of Hemp (Cannabis sativa L.) Cultivars for Total Delta-9 THC and Total CBD Using Gas Chromatography with Flame Ionization Detection. Plants 2024;13(13):519. 10.3390/plants13040519.

Berthold EC, Yang R, Sharma A et al. Regulatory sampling of industrial hemp plant samples (Cannabis sativa L.) using UPLC-MS/MS method for detection and quantification of twelve cannabinoids. J Cannabis Res 2020;2(2):42. 10.1186/s42238-020-00050-0.

Brighenti V, Pellati F, Steinbach M et al. Development of a new extraction technique and HPLC method for the analysis of non-psychoactive cannabinoids in fibre-type Cannabis sativa L. (hemp). J Pharm Biomed Anal 2017;143:228–36. 10.1016/j.jpba.2017.05.049.

Cao S, Kang S, Mao H et al. Defining molecular glues with a dual-nanobody cannabidiol sensor. Nat Commun 2022;13(13):815. 10.1038/s41467-022-28507-1.

Chand R, Mittal N, Srinivasan S et al. Upconverting nanoparticle clustering based rapid quantitative detection of tetrahydrocannabinol (THC) on lateral-flow immunoassay. Analyst 2021;146(146):574–80. 10.1039/D0AN01850C.

Chen C, Wang K, Wongso I et al. Postharvest blanching and drying of industrial hemp (Cannabis sativa L.) with infrared and hot air heating for enhanced processing efficiency and microbial inactivation. Dry Technol 2023;41(41):1454–68. 10.1080/07373937.2022.2159973.

De Prato L, Ansari O, Hardy GEStJ et al. The cannabinoid profile and growth of hemp (Cannabis sativa L.) is influenced by tropical daylengths and temperatures, genotype and nitrogen nutrition. Ind Crops Prod 2022;178:114605. 10.1016/j.indcrop.2022.114605.

Dilena E, Hunt I, Close DC. Optimal nitrogen rates and clonal effects on cannabinoid yields of medicinal cannabis. Sci Rep 2025;15(15):12341. 10.1038/s41598-025-96761-6.

Foight GW, Wang Z, Wei CT et al. Multi-input chemical control of protein dimerization for programming graded cellular responses. Nat Biotechnol 2019;37(37):1209–16. 10.1038/s41587-019-0242-8.

Garcia-de Heer L, Mieog J, Burn A et al. Weeding out variability: a proof-of-concept for producing uniform F1 hybrid Cannabis sativa L. using single-seed descent. Hortic Res 2026;13(13):uhag038. 10.1093/hr/uhag038.

Ghavami A, Labbé G, Brem J et al. Assay for drug discovery: Synthesis and testing of nitrocefin analogues for use as β-lactamase substrates. Anal Biochem 2015;486:75–7. 10.1016/j.ab.2015.06.032.

Guo Z, Smutok O, Ayva CE et al. Development of epistatic YES and AND protein logic gates and their assembly into signalling cascades. Nat Nanotechnol 2023;18(18):1327–34. 10.1038/s41565-023-01450-y.

Guo Z, Smutok O, Lee GR et al. Artificial allosteric protein switches with machine-learning-designed receptors. Nat Biotechnol 15 Apr. 2026:1–11. 10.1038/s41587-026-03081-9.

Harpaz D, Bernstein N, Namdar D et al. Portable biosensors for rapid on-site determination of cannabinoids in cannabis, a review. Biotechnol Adv 2022;61:108031. 10.1016/j.biotechadv.2022.108031.

Ii SLA, Pearson B, Kjelgren R et al. Response of essential oil hemp (Cannabis sativa L.) growth, biomass, and cannabinoid profiles to varying fertigation rates. PLOS ONE 2021;16(16):e0252985. 10.1371/journal.pone.0252985.

Jackson C, Anderson A, Alexandrov K. The present and the future of protein biosensor engineering. Curr Opin Struct Biol 2022;75:102424. 10.1016/j.sbi.2022.102424.

Kang S, Davidsen K, Gomez-Castillo L et al. COMBINES-CID: An Efficient Method for De Novo Engineering of Highly Specific Chemically Induced Protein Dimerization Systems. J Am Chem Soc 2019;141(141):10948–52. 10.1021/jacs.9b03522.

Li T, Bu J, Yang Y et al. A smartphone-assisted one-step bicolor colorimetric detection of glucose in neutral environment based on molecularly imprinted polymer nanozymes. Talanta 2024;267:125256. 10.1016/j.talanta.2023.125256.

Malík M, Praus L, Kuklina A et al. Cannabis yield and cannabinoid profile affected by plant nutrition and planting density. Ind Crops Prod 2025;223:120293. 10.1016/j.indcrop.2024.120293.

Miettinen K, Leelahakorn N, Almeida A et al. A GPCR-based yeast biosensor for biomedical, biotechnological, and point-of-use cannabinoid determination. Nat Commun 2022;13(13):3664. 10.1038/s41467-022-31357-6.

Milay L, Berman P, Shapira A et al. Metabolic Profiling of Cannabis Secondary Metabolites for Evaluation of Optimal Postharvest Storage Conditions. Front Plant Sci 2020;11. 10.3389/fpls.2020.583605.

Mulvihill CJ, Lutgens JD, Gollihar JD et al. A Humanized CB1R Yeast Biosensor Enables Facile Screening of Cannabinoid Compounds. Int J Mol Sci 2024;25(25):6060. 10.3390/ijms25116060.

Plouffe BD, Murthy SK. Fluorescence-based lateral flow assays for rapid oral fluid roadside detection of cannabis use. ELECTROPHORESIS 2017;38(3–4):501–6. 10.1002/elps.201600075.

Shaw WM, Zhang Y, Lu X et al. Screening microbially produced Δ9-tetrahydrocannabinol using a yeast biosensor workflow. Nat Commun 2022;13(13):5509. 10.1038/s41467-022-33207-x.

Stanton BZ, Chory EJ, Crabtree GR. Chemically induced proximity in biology and medicine. Science 2018;359(359):eaao5902. 10.1126/science.aao5902.

Tang J, Gao Z, Xu L et al. Smartphone-assisted colorimetric biosensor for the rapid visual detection of natural antioxidants in food samples. Food Chem 2025;462:141026. 10.1016/j.foodchem.2024.141026.

Wang JJ, Rodríguez Núñez JR, Maxwell EJ et al. Build Your Own Photometer: A Guided-Inquiry Experiment To Introduce Analytical Instrumentation. J Chem Educ 2015;93(93):166–71. 10.1021/acs.jchemed.5b00426.

Wohlwend J, Corso G, Passaro S et al. Boltz-1 Democratizing Biomolecular Interaction Modeling. Preprint, bioRxiv, 6 May 2025, 2024.11.19.624167. 10.1101/2024.11.19.624167.

