## Supplementary material for "Quantification of cannabidiolic acid in *Cannabis sativa* using a two-component allosteric protein biosensor"

**Supplementary Table S1.** Amino acids sequences of constructs used in this study. Where relevant specific components or linker elements have been highlighted using colour, bold or underlining.

| **Construct** | **Amino acid sequence** |
|---|---|
| **β-lactamase (TEM1)** | DHPETLVKVKDAEDQLGARVGYIELDLNSGKILESFRPEERFPMM<br>STFKVLLCGAVLSRVDAGQEQLGRRIHYSQNDLVEYSPVTEKHLT<br>DGMTVRELCSAAITMSDNTAANLLLTTIGGPKELTAFLHNMGDHV<br>TRLDRWEPELNEAIPNDERDTTMPVAMATTLRKLLTGELLTLASRQ<br>QLIDWMEADKVAGPLLRSALPAGWFIADKSGAGERGSRGIIAALG<br>PDGKPSRIVVIYTTGSQATMDERNRQIAEIGASLIKHW |
| **CaM-BLA 253** | HPETLVKVKDAEDQLGARVGYIELDLNSGKILESFRPEERFPMMST<br>FKVLLCGAVLSRVDAGQEQLGRRIHYSQNDLVEYSPVTEKHLTDG<br>MTVRELCSAAITMSDNTAANLLLTTIGGPKELTAFLHNMGDHVTRL<br>DRWEPELNEAIPNDERDTTMPVAMATTLRKLLTGELLTLASRQQLI<br>DWMEADKVAGPLLRSALPAGWFIADKSGAGERGSRGIIAALGPD<br>GSGGTEEQIAEFKEAFSLFDKDGDGTITTKELGTVMRSLGQNPTEA<br>ELQDMINEVDADGNGTIDFPEFLTMMARKMKDTDSEEEIREAFRV<br>FDKDGNGYISAAELRHVMTNLGEKLTDEEVDEMIREADIDGDGQ<br>VNYEEFVQMMTAGGSSGKPSRIVVIYTTGSQATMDERNRQIAEIG<br>ASLIKHW |
| **CaM-BLA 253-CA14** | DHPETLVKVKDAEDQLGARVGYIELDLNSGKILESFRPEERFPMM<br>STFKVLLCGAVLSRVDAGQEQLGRRIHYSQNDLVEYSPVTEKHLT<br>DGMTVRELCSAAITMSDNTAANLLLTTIGGPKELTAFLHNMGDHV<br>TRLDRWEPELNEAIPNDERDTTMPVAMATTLRKLLTGELLTLASRQ<br>QLIDWMEADKVAGPLLRSALPAGWFIADKSGAGERGSRGIIAALG<br>PDGSGGTEEQIAEFKEAFSLFDKDGDGTITTKELGTVMRSLGQNPT<br>EAELQDMINEVDADGNGTIDFPEFLTMMARKMKDTDSEEEIREAF<br>RVFDKDGNGYISAAELRHVMTNLGEKLTDEEVDEMIREADIDGDG<br>QVNYEEFVQMMTAGGSSGKPSRIVVIYTTGSQATMDERNRQIAEI<br>GASLIKHWGGSGSGGSGSGSGSSGSEVQLQASGGGFVQPGGS<br>LRLSCAASGSTSRQYDMGWFRQAPGKEREFVSAISSNQDQPPYY<br>ADSVKGRFTISRDNSKNTVYLQMNSLRAEDTATYYCAFKQHHAN<br>GAYWGQGTQVTVSSKLAAALEHHHHHH |
| **DB21-CaM BP** | DGHHHHHHGSGSEVQLQASGGGFVQPGGSLRLSCAASGTTYG<br>QTNMGWFRQAPGKEREFVSAISGLQGRDLYYADSVKGRFTISRD<br>NSKNTVYLQMNSLRAEDTATYYCAFHDFLRMWEYWGQGTQVTV<br>SSGGGSGSSGGSGGSGGGKRRWKKNFIAVSAANR |

| | |
|---|---|
| **CaM-BLA 253-DB21** | DHPETLVKVKDAEDQLGARVGYIELDLNSGKILESFRPEERFPMM<br>STFKVLLCGAVLSRVDAGQEQLGRRIHYSQNDLVEYSPVTEKHLT<br>DGMTVRELCSAAITMSDNTAANLLLTTIGGPKELTAFLHNMGDHV<br>TRLDRWEPELNEAIPNDERDTTMPVAMATTLRKLLTGELLTLASRQ<br>QLIDWMEADKVAGPLLRSALPAGWFIADKSGAGERGSRGIIAALG<br>PDGSGGTEEQIAEFKEAFSLFDKDGDGTITTKELGTVMRSLGQNPT<br>EAELQDMINEVDADGNGTIDFPEFLTMMARKMKDTDSEEEIREAF<br>RVFDKDGNGYISAAELRHVMTNLGEKLTDEEVDEMIREADIDGDG<br>QVNYEEFVQMMTAGGSSGKPSRIVVIYTTGSQATMDERNRQIAEI<br>GASLIKHWGGSGSGGSGSGSGSSGSEVQLQASGGGFVQPGGS<br>LRLSCAASGTTYGQTNMGWFRQAPGKEREFVSAISGLQGRDLYY<br>ADSVKGRFTISRDNSKNTVYLQMNSLRAEDTATYYCAFHDFLRM<br>WEYWGQGTQVTVSSKLAAALEHHHHHH |
| **CA14-CaM BP** | DGHHHHHHGSGSEVQLQASGGGFVQPGGSLRLSCAASGSTSR<br>QYDMGWFRQAPGKEREFVSAISSNQDQPPYYADSVKGRFTISRD<br>NSKNTVYLQMNSLRAEDTATYYCAFKQHHANGAYWGQGTQVTV<br>SSGGGSGSSGGSGGSGGGKRRWKKNFIAVSAANR |
| **CA14$_{D34S}$-CaM BP** | DGHHHHHHGSGSEVQLQASGGGFVQPGGSLRLSCAASGSTSR<br>QYSMGWFRQAPGKEREFVSAISSNQDQPPYYADSVKGRFTISRD<br>NSKNTVYLQMNSLRAEDTATYYCAFKQHHANGAYWGQGTQVTV<br>SSGGGSGSSGGSGGSGGGKRRWKKNFIAVSAANR |

**Supplementary Figure S1**

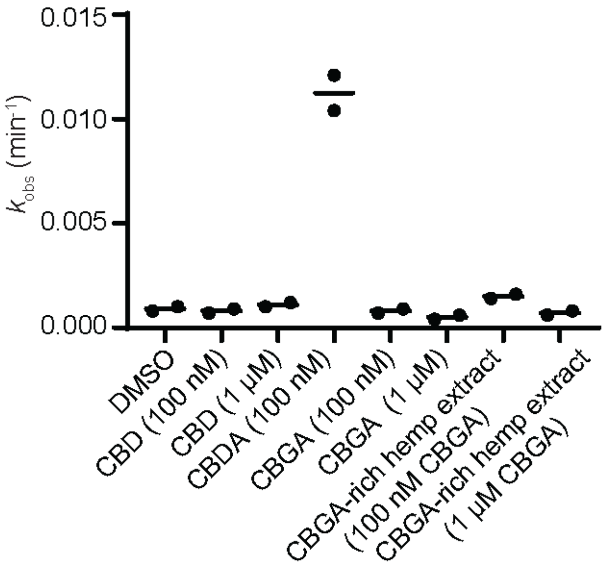


**Supplementary Figure S1 Biosensor response to hemp extract.** Rate of UW154 hydrolysis (ΔA520 nm), reported as observed rate constant ($k_{obs}$, min$^{-1}$), in response to different concentrations of CBD, CBDA, CBGA, and CBGA-rich hemp extract diluted to a final concentration of 100 nM or 1 μM CBGA. CBDA biosensor (10 nM CaM-BLA-DB21 and 100 nM $CA14_{D34S}$-CaM BP) was incubated with 50 μM UW154 and different cannabinoid-containing substrates at 25 °C.

**Supplementary Figure S2**

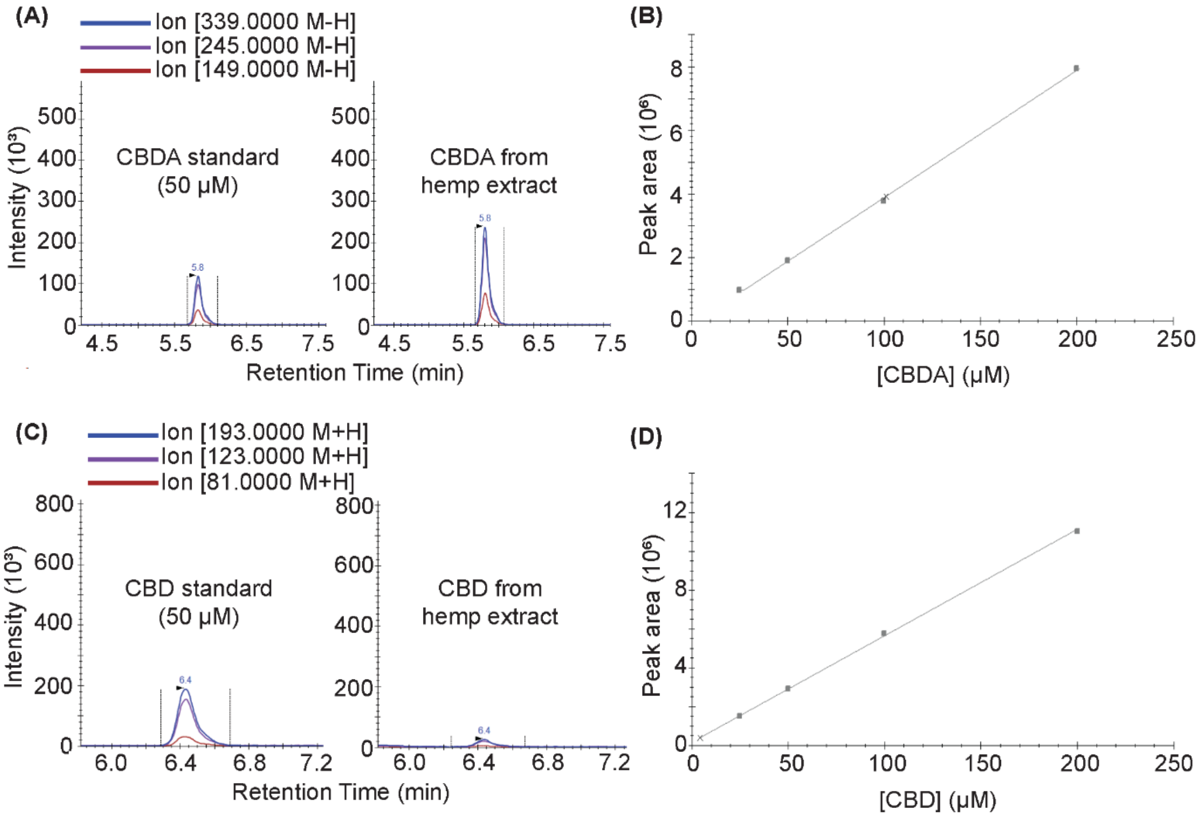


**Supplementary Figure S2 Quantification of CBDA and CBD in hemp biomass extracts.** **(A)** Sample chromatograms from LC-MS/MS (negative ion mode) analysis of CBDA. Target ion peaks are shown in the relevant CBDA retention time window for a CBDA analytical reference standard (50 μM) and a redissolved extract from CBDA-rich hemp (prior to further dilution. **(B)** Standard curve of CBDA (peak area of target ion 339). **(C)** Sample chromatograms from LC-MS/MS (positive ion mode) analysis of CBD. Target ion peaks are shown in the relevant CBD retention time window for a CBD analytical reference standard (50 μM) and a redissolved extract from CBDA-rich hemp (prior to further dilution). **(D)** Standard curve of CBD (peak area of target ion 193).
